# Counterfactual Explanations for Graph Neural Networks in Patient Outcome Prediction

**DOI:** 10.64898/2026.05.18.725906

**Authors:** Nikolaos Chaidos, Angeliki Dimitriou, Hari Calzi, Elena Casiraghi, Giorgos Stamou, Giorgio Valentini

## Abstract

Counterfactual Explanation (CE) algorithms have been successfully applied to uncover the main factors driving computational diagnostic and prognostic predictions on tabular medical data. Recently, a new Network Medicine paradigm has been introduced for patient diagnosis and prognosis using Patient Similarity Networks (PSNs), i.e. graphs where patients are represented as nodes and their clinical and biomolecular similarities as edges. In this context, graph-based algorithms, including Graph Neural Networks (GNNs), can provide predictions using not only individual patient features but also their relations within a network of clinically and biomolecularly similar individuals. In this work, we propose the first CE algorithm tailored to explain diagnostic and prognostic predictions within PSNs. Alongside a contrastive GNN backbone, we introduce a versatile, model-agnostic counterfactual search method compatible with any underlying classifier. Preliminary results on synthetic data and on a cohort of patients affected by the Alzheimer’s disease show that our algorithm is competitive both with seminal tabular based CE algorithms and GNNExplainer, a well-established method for explaining graph-based classification tasks.

## 1 Introduction

The explainability of AI model predictions remains a critical bottleneck in the clinical adoption of healthcare applications [14]. Initially, the interpretability of black-box models on medical data relied on local feature attribution methods, such as LIME [17] and SHAP [12]. While these techniques highlight which features contribute to a specific prediction, they fail to explicitly guide clinicians on how to achieve a different clinical outcome, e.g. by identifying which factors may be responsible for changing an unfavorable prognosis into a favorable one, based on a patient’s individual characteristics.

To bridge this gap, Wachter et al. [21] introduced counterfactual explanations as an optimization problem, seeking the minimal perturbation to input features required to flip a prediction. State-of-the-art methods have since evolved to address the limitations of these early approaches. Frameworks like DiCE [13] generate diverse options, giving clinicians a range of potential alternatives, while recent advancements, such as FACE [15], enforce actionability and causal constraints. A major limitation of these tabular methods is their fundamental assumption that instances (patients) are entirely unrelated and independent.

Recent systems biology approaches represent patients and their relationships as graphs (i.e. Patient Similarity Networks - PSN), where nodes are patients and edges correspond to their integrated biomolecular and clinical similarities [6]. When PSNs are processed with graph-based models, predictions can exploit not only individual patient features but also each patient’s relational context within a network of clinically and biomolecularly similar individuals [5].

In the broader domain of Graph Neural Networks (GNNs), explainability has similarly evolved from factual importance (e.g., GNNExplainer [25]) to generating actionable counterfactuals (e.g., CF-GNNExplainer [11] and CF^2^ [19]). Yet, despite these advancements, a significant gap remains in biomedical applications.

Even if some attempts to design explainable prediction methods in biomedical graphs exist, they are limited to brain networks [1] or Electronic Medical Records [24]. Furthermore, as outlined by a recent review [16], existing counterfactual GNN methods primarily focus on graph classification tasks.

To our knowledge, no explainable methods have been explicitly designed to generate counterfactuals for vertex classification in PSNs; the exact task required for personalized diagnosis and outcome prediction.

o address this unexplored problem, this paper presents a novel *Counterfactual Explanation Graph Neural Network (CE-GNN)* framework explicitly designed for PSNs. This paper establishes a new approach to personalized patient prognosis or outcome prediction by integrating multi-omics data, PSN construction, and a novel GNN-based counterfactual prediction algorithm. Because this represents a first exploration into PSN-specific counterfactuals, the proposed framework is inherently preliminary. Consequently, we detail both the merits and the limitations of this current version to guide future research, ultimately demonstrating the feasibility and transformative potential of counterfactual network medicine in precision healthcare.

## 2 Method

Our method couples the integration of multi-modal bio-medical data with a novel explanation framework. Data are first structured into graphs using the Similarity Network Fusion (SNF) algorithm [22] (Section 2.1), with the potential to utilize the miss-SNF algorithm to manage completely missing data in subsets of patients [4]. Once the network is constructed, we apply our newly proposed CE-GNN algorithm to explicitly provide explainable predictions for personalized patient outcomes (Section 2.2).

### 2.1 Patient similarity network integration through SNF

To construct a unified Patient Similarity Network (PSN) from multiple data views, we utilize the SNF algorithm [22]. SNF operates by first using a scaled exponential similarity kernel to compute a pairwise similarity matrix between the biomolecular profiles of individuals for each independent data source. To capture both the broad relationships and the fine-grained local topology, SNF derives a global similarity matrix (representing overall patient relationships) and a local similarity matrix (restricted to a patient’s *k*-nearest neighbors) for each modality. An iterative cross-diffusion process is then applied, where the local similarities of one modality guide the update of the global similarities in the others. This message-passing process continues until convergence, resulting in a single, integrated network *G* that robustly synthesizes information across all original views. For the complete mathematical formulation of the underlying kernels and update rules, we refer readers to [22].

### 2.2 The CE-GNN Framework

The output of the data integration phase (Section 2.1) is a unified PSN graph *G* = (*V, E*), where nodes *v* ∈ *V* correspond to patients and edges *e* ∈ *E* denote their bio-molecular similarities computed by SNF. We define a labeling function *ψ* : *V* → {*C*_1_, *C*_2_, …, *C*_*m*_} to assign each patient to one of *m* discrete classes. In a standard binary classification task (*m* = 2), these classes typically represent opposing clinical outcomes, such as poor (*C*_1_) versus good (*C*_2_) prognosis.

#### Contrastive Learning for PSN Node Representation

Each patient node in the PSN is initially attributed with a feature vector comprising its corresponding clinical and multi-omics data. To learn meaningful embeddings from these features and the graph structure, we employ a Siamese GNN, inspired by counterfactual methodologies such as those in [3] and [2]. This framework can be instantiated using standard message-passing architectures, such as Graph Convolutional Networks (GCN) [10] or Graph Attention Networks (GAT) [20]. To optimize the network, we train it to minimize the *contrastive loss* function:

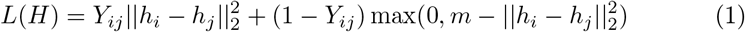

where: **i)** *h*_*i*_, *h*_*j*_ ∈ ℝ^*d*^ denote the *d*-dimensional node embeddings for patients *v*_*i*_ and *v*_*j*_ computed by the GNN, corresponding to rows within the hidden representation matrix *H* ∈ ℝ^*n×d*^, where *n* is the number of patients, **ii)** *Y*_*ij*_ is a binary indicator variable defined as *Y*_*ij*_ = 1 if *ψ*(*v*_*i*_) = *ψ*(*v*_*j*_) (i.e., the patients belong to the same clinical class), and *Y*_*ij*_ = 0 otherwise, and **iii)** *m* represents a predefined margin parameter, ensuring that dissimilar patient pairs (*h*_*i*_, *h*_*j*_) that are separated by a Euclidean distance greater than *m* do not incur a loss penalty.

Intuitively, this contrastive objective forces the GNN to learn an embedding space where patients sharing the same clinical diagnosis or outcome are clustered closely together, while patients with differing outcomes are pushed apart. Once this discriminative embedding space is established, final class predictions are obtained by training a simple linear classifier on top of these expressive GNN representations using a standard cross-entropy loss.

### Ranking and Counterfactual Retrieval

For each query node *v*_*i*_ with hidden representation *h*_*i*_, we retrieve a counterfactual example, *v*_*ci*_, by selecting the most similar node *v*_*j*_ whose predicted class differs from that of *v*_*i*_, i.e., *ψ*(*v*_*j*_) ≠*ψ*(*v*_*i*_). Similarity is measured using cosine similarity in the embedding space. Formally, the retrieved counterfactual node is defined as:

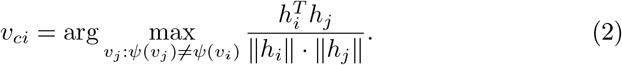

The corresponding counterfactual hidden representation is denoted by *h*_*ci*_, i.e., the embedding associated with the retrieved node *v*_*ci*_. In the following, *f*^*ci*^ denotes the feature vector associated with *v*_*ci*_, while *f*^*i*^ denotes the feature vector associated with the original query node *v*_*i*_.

#### Counterfactual Feature Extraction via Iterative Swapping

While the overall CE-GNN framework integrates the entire data-to-prediction pipeline, this final stage isolates the exact explanation. We define an explanation as a minimal, interpretable set of feature-level edits required to conceptually transform patient *v*_*i*_ into *v*_*ci*_, thereby changing the model’s prediction. Ideally, these edits represent plausible, actionable interventions in a real-world medical context.

Given the feature vectors 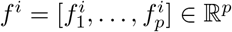 and 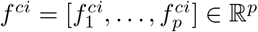, the general intuitive idea is to progressively swap a growing subset of *f*^*i*^ of patient *v*_*i*_ with the features *f*^*ci*^ of the most similar “counterfactual patient” *v*_*ci*_ until the trained GNN changes its prediction. The resulting subset of features represents the *counterfactual explanation* for that patient.

To achieve this, we apply the following sequential extraction process for each patient *v*_*i*_:

- *Feature Normalization:* To ensure a fair comparison of magnitudes across different biological or clinical modalities, all features are first standardized (e.g., via z-score or softmax normalization).
- *Feature Ranking Strategies:* To determine the optimal order in which features should be swapped, we rank them based on their likelihood to drive a change in prediction. Our framework implements three distinct ranking mechanisms: i) Absolute Difference: A baseline approach computing the absolute element-wise divergence (| *f*^*i*^ *™ f*^*ci*^ |) between the target patient and their counterfactual, assuming the largest raw differences are the primary drivers, ii) Statistical Significance: A data-driven approach computing the ANOVA F-values to assess the correlation and importance of each individual feature against the clinical outcome, iii) Model-Aware Importance: A predictive approach that extracts intrinsic feature importance scores directly from an auxiliary trained Random Forest classifier to guide the substitution order based on learned predictive power.
- *Iterative Perturbation:* Following the ranked order computed in the previous step, we cumulatively assign one by one the specific feature values of the counterfactual patient *v*_*ci*_ to the feature vector of the target patient *v*_*i*_.
- *Explanation Extraction:* After every individual feature swap, the modified feature vector is evaluated by the trained GNN. The iterative loop breaks when the GNN predicts a new class label. The precise subset of features responsible for inducing this label switch is recorded as the counterfactual explanation *s*_*i*_. If the entire feature vector is swapped without triggering a change in the prediction, *s*_*i*_ is set to NULL.

Ultimately, this process returns a list of counterfactual explanations for patients in the network, highlighting the most critical personalized factors underlying their specific diagnosis/prognosis.

#### Instance and class-level explanations

The proposed approach can be considered an **instance-level (local) explanation method**, since the extracted feature subset *s*_*i*_ explicitly highlights the personalized factors driving the the switching of the prediction for each individual patient.

We can further derive **class-level (global) explanations** *S*(*C*_*k*_) by combining and stratifying by classes 𝒞 = {*C*_1_, *C*_2_, …, *C*_*m*_} the individual counterfactual features *s*_*i*_ that led to a switching of the prediction:

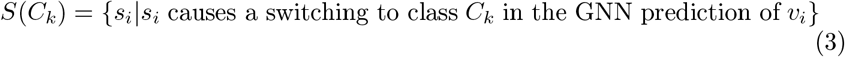

Then for each *C*_*k*_ ∈ 𝒞 we can select the most important common counterfactual features by intersection, union or counting. For instance in the case of the counting approach we have:

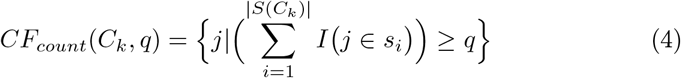

where *j* are the indices of the selected counterfactual features, *q* is a parameter such that 1 ≤ *q* ≤ *n*, and *I*(*x*) is an indicator function. Eq. 4 requires that a feature can be included in *CF*_*count*_(*C*_*k*_, *q*) if selected at least *q* times.

## 3 Experimental Evaluation

### 3.1 Datasets

Our work focuses on the design and application of CE algorithms to networked bio-medical data. However, because real-world clinical datasets lack a definitive ground truth for feature-level explanations, it is difficult to objectively evaluate explanation accuracy. Therefore, to validate whether the CE-GNN can successfully identify and extract significant counterfactual features, we apply our algorithm to synthetic data where the ground-truth features are known by design. It should be noted that the proposed CE-GNN pipeline and methodology remains exactly the same for both synthetic and real-world datasets.

#### Synthetic networked data

We first generated tabular synthetic data representing different views of the examples belonging to two classes with a pre-defined subset of discriminative features. The synthetic dataset comprises *n* = 300 samples belonging to two distinct classes 𝒞 ∈ {0, 1} and *p* = 150 features, partitioned into three distinct views to simulate the heterogeneity of multi-omics data sources used in PSN construction. Features of Class 0 examples are distributed according to a 0-centered normal distribution 𝒩 (0, *σ*) having different standard deviations *σ* and Class 1 examples are characterized by specific stochastic patterns restricted to subsets of features for each specific view (Fig. 1).

**Fig. 1.**
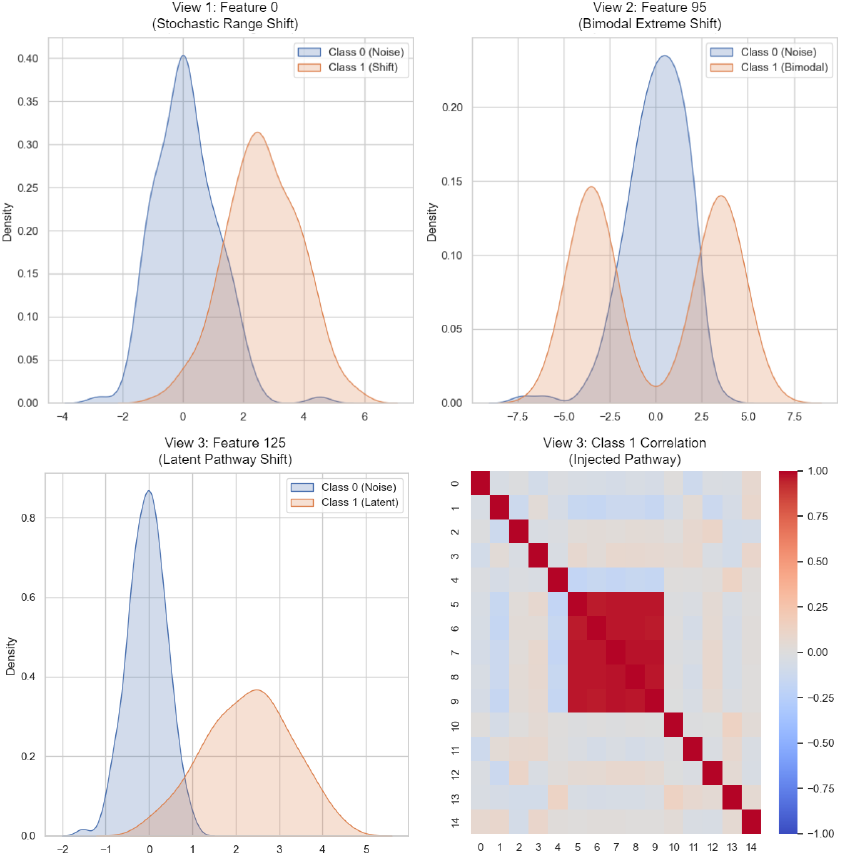
Profiles of synthetic features. Class 1 introduces discriminative signals across three views: a stochastic range shift (View 1, top-left), a bimodal toggle (View 2, top-right), and a latent co-expression pathway (View 3, bottom-left). The figure at the bottom right represents the correlation between Class 1 examples in View 3.

#### View 1 (Intermediate *σ* = 1.0)

Features *j* ∈ [0, 4] exhibit a **Stochastic Range Shift**. For Class 1, values are drawn from 𝒩 (0, 1) + 𝒰(1.5, 4.0), simulating a standard high-throughput modality where a variable positive shift is the primary driver. **View 2 (High-Variance** *σ* = 1.5): Features *j* ∈ [95, 99] utilize a **Bimodal Toggle**. Class 1 instances are randomly assigned to either high positive or high negative states (± 3.5). **View 3 (Low-Variance** *σ* = 0.5): Features *j* ∈ [125, 129] follow a **Latent Co-expression** pattern. A single latent biological driver controls this 5-feature block, testing the explainer’s ability to identify correlated functional pathways in relatively clean data.

#### The ROSMAP dataset

To evaluate the framework in a real-world clinical context, we utilize a multi-omics Alzheimer’s disease dataset, originally curated to benchmark the MOGONET architecture [23]. The cohort comprises 351 patients (182 individuals diagnosed with Alzheimer’s, 169 controls). The dataset encompasses three distinct biological modalities: mRNA expression, miRNA expression, and DNA methylation data. To ensure consistency with established baselines, we utilize the data exactly as pre-processed in [23], which isolates the 200 most informative features for each respective omics modality.

### 3.2 Metrics and Baselines

We assess the performance of the explanation methods across two primary dimensions: their ability to accurately identify the underlying drivers of a prediction and the structural quality of the generated counterfactuals. For the first dimension, we compare the predicted subset of important features to the ground-truth important features. Since ground-truth features are known by design only for the synthetic dataset, this evaluation is restricted to the synthetic experiments. We frame this as a binary classification task, where a feature is either correctly or incorrectly identified as important, and utilize *Precision, Recall, Accuracy*, and *Macro-F1* metrics.

For the second dimension, to validate the counterfactual generation aspect of the models, we measure *Validity, Fidelity*, and *Sparsity*, according to the framework proposed in [16]. Validity (or the flip rate percentage) assesses the proportion of generated counterfactuals that successfully alter the initial prediction. Fidelity measures how faithful the explanations are to the oracle (the trained classifier model) considering their correctness. Sparsity measures the conciseness of the explanation by quantifying the relative reduction in feature space. The formal definitions for these metrics are:

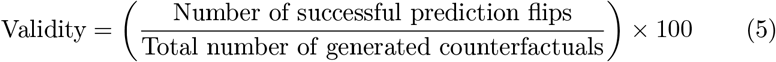

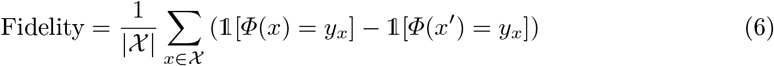

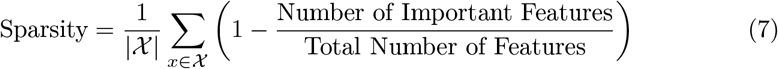

where 𝟙 [*Φ*(*x*) = *y*_*x*_] is equal to 1 if the classifier’s predicted class for the original sample *x* is its true label *y*_*x*_, and 𝟙 [*Φ*(*x*′) = *y*_*x*_] is equal to 1 if the classifier’s predicted class for the generated counterfactual *x*′ is the label *y*_*x*_. Fidelity ranges from −1 to 1, where a value of 1 indicates a perfect classifier and counterfactual generator. Validity and Sparsity range from 0 to 1, where higher values indicate better performance (note that all metrics are scaled to percentages in our reported tables).

To comprehensively evaluate our approach, we compare our method against three established explainability baselines: DiCE [13], SingleCF (Wachter et al. [21]), and GNNExplainer [25]. This selection allows us to benchmark our framework against both traditional counterfactual tabular-based approaches (DiCE, SingleCF) as well as a prominent factual graph-based approach (GNNExplainer). Because Validity and Fidelity explicitly evaluate counterfactual transitions, GN-NExplainer is not evaluated for these two metrics since it is purely a factual method. For each method, we extract a final subset of predicted important features to properly compute and compare each metric. Our proposed extraction process via iterative swapping is detailed in Section 2.2. For the tabular counterfactual baselines (DiCE and SingleCF), we isolate the important features by selecting those that exhibit an absolute difference of at least 0.05 between the original patient vector and the counterfactual vector, a standard threshold chosen to account for the fact that all features undergo z-score normalization prior to extraction. Finally, for GNNExplainer, we take the continuous importance scores it outputs for all features and optimally cluster them into two distinct groups using the Jenks natural breaks optimization method [9]; the cluster containing the higher importance scores is subsequently treated as the final discrete subset of important features. Finally for our proposed method, we implement all three different feature ranking methods (**Abs**olute-Difference, **St**atistical and **R**andom-**F**orest) mentioned in Section 2.2.

#### Implementation Details

The the GNN backbone of our method for both datasets was instantiated as a 3-layer Graph Convolutional Network (GCN) with 256 hidden channels and a dropout rate of 0.3. For the synthetic dataset, the GCN utilized LayerNorm and residual connections, while for the ROSMAP dataset, it utilized BatchNorm without residuals. We set the contrastive loss margin to 2.0, learning rate to 0.001 and trained the backbone for 400 epochs.

### 3.3 Results

The underlying base classifier (our contrastive GCN) achieved a robust test set accuracy of 96.67%, providing a strong predictive foundation for explanation extraction. The evaluation on the synthetic dataset (Table 1) demonstrates that the proposed CE-GNN framework, particularly when utilizing Random Forest (RF) feature ranking, effectively isolates the key factors behind a prediction.Traditional tabular counterfactual baselines, such as SingleCF and DiCE, struggle with identifying the designed ground-truth features, achieving a precision of roughly 10% and a macro-F1 score of approximately 17%. While these tabular baselines yield higher recall by selecting broader feature sets, CE-GNN (RF) provides a superior balance of precise feature isolation, reaching 60.59% precision and 46.87% macro-F1. Additionally, CE-GNN (RF) maintains a highly competitive accuracy of 86.73%, trailing only slightly behind the factual baseline GNNExplainer. Furthermore, while the factual baseline GNNExplainer achieves the highest sparsity (92.30%), CE-GNN (RF) not only achieves the best feature-recovery performance, it also maintains a competitive high sparsity (89.09%), perfect validity (100%), and strong fidelity (96.67%).

**Table 1.**
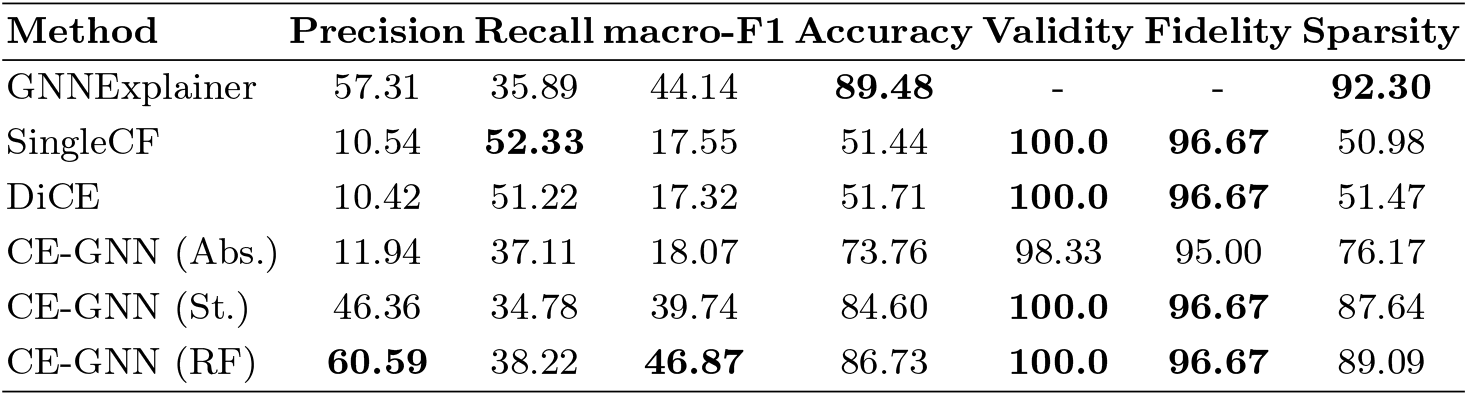
Comparison across methods on the Synthetic Dataset, including evaluation according to the pre-defined ground-truth important features.

| Method | Precision | Recall | macro-F1 | Accuracy | Validity | Fidelity | Sparsity |
| --- | --- | --- | --- | --- | --- | --- | --- |
| GNNExplainer | 57.31 | 35.89 | 44.14 | **89.48** | - | - | **92.30** |
| SingleCF | 10.54 | **52.33** | 17.55 | 51.44 | **100.0** | **96.67** | 50.98 |
| DiCE | 10.42 | 51.22 | 17.32 | 51.71 | **100.0** | **96.67** | 51.47 |
| CE-GNN (Abs.) | 11.94 | 37.11 | 18.07 | 73.76 | 98.33 | 95.00 | 76.17 |
| CE-GNN (St.) | 46.36 | 34.78 | 39.74 | 84.60 | **100.0** | **96.67** | 87.64 |
| CE-GNN (RF) | **60.59** | 38.22 | **46.87** | 86.73 | **100.0** | **96.67** | 89.09 |

For the ROSMAP dataset, the base GCN classifier achieved a test set accuracy of 70.75%. Transitioning to this real-world clinical context, the results (Table 2) highlight the CE-GNN framework’s advantage in producing highly concise explanations. Baseline tabular methods like SingleCF and DiCE achieve a 100% validity rate and 70.75% fidelity; however, they rely on relatively dense feature sets to flip the prediction, as evidenced by their lower sparsity scores of 49.69% and 51.24%, respectively. Consequently, these tabular baselines perturb roughly half of the 600 available features to generate a single counterfactual instance, yielding explanations that are too broad to be practically useful. Conversely, the CE-GNN variants maintain strong predictive performance while drastically reducing the feature space required for a counterfactual transition. Specifically, CE-GNN (Abs.) and CE-GNN (RF) achieve high sparsity scores of 85.75% and 77.95%, respectively, while retaining robust validity rates of 91.51% and 90.57%.

**Table 2.** Comparison across methods on ROSMAP dataset.

| Method | Validity | Fidelity | Sparsity |
| --- | --- | --- | --- |
| GNNExplainer | - | - | 80.17 |
| Wachter | **100.0** | **70.75** | 49.69 |
| DiCE | **100.0** | **70.75** | 51.24 |
| CE-GNN (Abs.) | 91.51 | 62.26 | **85.75** |
| CE-GNN (St.) | 85.85 | 56.60 | 75.69 |
| CE-GNN (RF) | 90.57 | 61.32 | 77.95 |

Biomedical literature confirms that most of the counterfactual genomic and epigenomic features selected by CE-GNN, with Equations 3 and 4 (approximately 30 when Random Forests are used to rank the features), are associated with Alzheimer’s disease (AD). For example, the miRNA hsa-miR-34a is strongly overexpressed in specific AD brain regions [18], whereas hsa-miR-132 has been repeatedly reported as dysregulated or downregulated in AD [26]. In addition, miR-129-5p has been identified as a biomarker for pathology and cognitive decline in Alzheimer’s disease [7], while activation of the gene GPR120 has been shown to inhibit amyloid pathology associated with AD [8].

## 4 Conclusions

Counterfactual explanations of medical decisions can individuate actionable factors that may change the outcome of a patient; for example, shifting a prognosis from poor to good. While these methods have been largely applied to predictive computational models trained on tabular data across different medical domains, to our knowledge, no CE methods have yet been applied in the relational framework of Patient Similarity Networks. Our work represents a first attempt to apply CE strategies to explain GNN predictions for disease diagnosis, combining the discovery of counterfactual features with the integration of multi-omics data. Our results show that our proposed method is competitive with other established approaches and can provide sparse counterfactual features.

It is important to note that the algorithm cannot currently generate counterfactuals for all patients in the ROSMAP dataset, leading to a validity of less than 100% (Table 2). This limitation occurs because our current algorithm only modifies the features of the patients associated with the PSN nodes, whereas GNN predictions also strongly depend on the topology the graph. For this reason, by keeping the topology of the graph unchanged, modifying node features alone is insufficient to change the prediction of the GNN for certain patients. Hence a natural extension of our approach is an algorithm that is capable of changing both the node features and the graph edges in order to improve the validity of our method. Moreover, further experiments across diverse datasets are necessary to confirm these preliminary findings.

Nevertheless, to the best of our knowledge, our approach is the first application of a CE approach in the context of graph-based diagnostic predictions, paving the way for novel applications of CE algorithms within Network Medicine.

## Acknowledgments

This work was partially funded by Piano di Sviluppo di Ricerca (PSR2025) - Università degli Studi di Milano.

Part of this work was funded by the National Plan for NRRP Complementary Investments (PNC) in the call for the funding of research initiatives for technologies and innovative trajectories in the health—project n. PNC0000003—AdvaNced Technologies for Human-centrEd Medicine (project acronym: ANTHEM).

This work was also supported by the Hellenic Foundation for Research and Innovation (HFRI) under the 5th Call for HFRI PhD Fellowships (Fellowship Number 19268).

Computational resources were provided by the INDACO Core facility (University of Milan).

